# Delineation of Single-cell Altas Provides New Insights for Development of Coronary Artery Lesions in Kawasaki Disease: Bad and Good Signaling Molecules

**DOI:** 10.1101/2024.11.13.623351

**Authors:** Qiuping Lin, Xin Lv, Qingzhu Qiu, Lianni Mei, Liqin Chen, Sirui Song, Wei Liu, Xunwei Jiang, Min Huang, Libing Shen, Tingting Xiao, Lijian Xie

**Author notes:** Correspondence: Libing Shen,; Tingting Xiao,; Lijian Xie. These authors contributed equally (Qiuping Lin, Xin Lv, Qingzhu Qiu).

## Abstract

**Background:** Kawasaki Disease (KD) is a vasculitis syndrome featured with a high and persistent fever and mainly affects children under 5 years of age. It is the leading cause of acquired heart disease in the form of coronary artery lesions (CALs) for children in developed countries. Most KD can be relieved with the high-dosage Intravenous immunoglobulin (IVIG) therapy, but a small proportion develop CALs after IVIG treatment.

**Methods:** We performed the single-cell RNA sequencing for the peripheral blood mononuclear cells (PBMCs) from three KD non-CAL patients whose samples were acquired before and after IVIG treatment and three KD CAL patients whose samples were acquired only after IVIG treatment. Cell-to-cell communication patterns were also analyzed in KD CAL and non-CAL patients

**Results:** Overall cell expression feature analyses show immunoglobulin and adaptive immunity related genes are upregulated in KD CAL patients while B cell activation related genes are downregulated in them. Pseudo-time analyses demonstrate that both KD non-CAL patients before treatment and KD CAL patients after treatment have a dysregulated B cell developmental trajectory while the former has a mixed T and B lineage and the latter has a mixed monocyte and B lineage. The early elevated expression of SPI1 could partly explain the dysregulated B cell development in KD CAL patients. Cell communication analyses propose a disorder cell communication pattern in KD non-CAL patients before treatment and some persistent bad cell-to-cell signals in KD CAL patients after treatment. There are four signaling molecules, APP, CCL, and MCH-II, whose expression is significantly increased in the CD14 and CD16 monocytes of KD CAL patients, where the expression of RESISTIN is significantly increased in those KD non-CAL patients.

**Conclusions:** Our results suggest that APP, CCL, and MCH-II might be the bad signals for indicating the possible development of CAL while RESISTIN is a good one for protecting from CAL.

## Introduction

Kawasaki disease (KD) is a form of small and medium-sized vasculitis featured with high fever and its etiology is unknown until now. KD usually occurs in children under the age of 5 years, causing coronary artery dilatation or aneurysm, and it is the most common cause of acquired heart disease in children in developed countries^1^. If KD was left untreated, the incidence of coronary artery lesions (CALs) is about 20% to 25%^2^. Intravenous immunoglobulin (IVIG) is the standard treatment for KD, which reduces the risk of CALs to 5%^3^.

The pathological process of CALs is not clear at present. Studies figured out that coronary vasculitis begins 6–8 days after the onset of KD and then the inflammation rapidly invades all layers of the artery^4^. Coronary artery aneurysms (CAAs) have been reported to regress in size after 4–8 weeks of an acute episode, but it may take several years for coronary artery diameters to return to normal^1^. Orenstein et al identified three interrelated pathologic processes in KD with CALs, including necrotizing arteritis, subacute chronic arteritis, and luminal myofibroblast proliferation^5^.The endothelium, located in the inner surface of coronary arteries, serves as the interface between the circulation and the vascular media or adventitia, so endothelial dysfunction (ED) is believed to contribute to the development of CALs^6,7^. Studies suggest that CALs formation in KD are related to various genetic factors, such as HLA-E, HLA-B, CASP3, ITPKC, and etc.^8–10^. Inflammatory cell activation, pro-inflammatory cytokines production, the innate immune system and the adaptive immune system are involved in vasculitis formation^11^. But the mechanism of CALs formation still remains unknown, which brings challenges to the prevention and treatment of CALs.

Unlike traditional bulk RNA sequencing methods, single-cell RNA sequencing (scRNA-seq) allows for the interrogation of individual cells, revealing distinct transcriptional states and facilitating the identification of rare cell populations^12^. In conjunction with the capabilities of scRNA-seq, the exploration of cell-to-cell communication has gained significant attention^13^. By analyzing cell communication patterns to infer and visualize signaling pathways between cells^14^, Cell-to-cell communication can elucidate how different cell types interact in a dynamic microenvironment. This understanding of cell communication is crucial for deciphering complex biological processes, such as immune responses and tissue homeostasis^15^. scRNA-seq combined with cell communication analysis were applied for pancreatic, and breast cancers, elucidating complex tumor microenvironments and identifying novel therapeutic approaches^16–18^. What’s more, these methodologies were applied for Immune-mediated diseases, delineating disease-specific ligand-receptor networks and shed light on shared immune responses across tissues and diseases^19^.

So far, the studies investigating KD with CALs by analyzing cellular communication at the single cell level are still lacking. Here we performed single-cell transcriptomic sequencing on six children with KD. Total nine scRNA-seq samples were acquired and three of them developed CALs after IVIG treatment. Cell-to-cell communication was analyzed among different sample groups. Our study aims to explore cellular panorama and communication networks in KD, ultimately contributing to the understanding of its etiology and potential therapeutic strategies.

## Results

### Clinical information of KD CAL and non-CAL patients

All KD patients with CALs (KL) of the study were recruited from the Shanghai Children’s Hospital between August 2023 and April 2024. The diagnosis of complete and incomplete KD is according to 2017 American Heart Association (AHA) guidelines ^1^. The maximum internal diameter of the coronary arteries was obtained through echocardiography. These measurements were converted to Z scores using a model derived from the Kabayashi method^20^. The CALs were defined according to the 2017 AHA Guidelines^1^. Blood samples were collected from all patients after confirmed diagnoses of complete KD and after IVIG treatment. The non-CAL KD patients were from our previous research (NA1022839) (Table 1). Informed consent was obtained from the guardians of the children. All manipulations were approved by the Ethics Committee of Shanghai Children’s Hospital (IRB number: 2022R121)

**Table 1.** Demographics for Kawasaki Disease Patients.

|  | KD1 | KD2 | KD3 | KL1 | KL2 | KL3 |
| --- | --- | --- | --- | --- | --- | --- |
| Age(years) | 2.1 | 5.6 | 5.4 | 1.5 | 1.0 | 2.0 |
| Sex(M/F) | M | M | F | F | F | M |
| Fever(days) | 5 | 5 | 5 | 14 | 5 | 5 |
| Diagnosis | cKD | cKD | iKD | cKD | cKD | cKD |
| IVIG responsive | Yes | Yes | Yes | Yes | Yes | Yes |
| Z score of the blood draw time | 1.02 | 1.84 | 0.71 | 4.11 | 4.25 | 2.66 |

### Delineation of single-cell transcription atlas of PBMCs in KD CAL and non-CAL patients

To delineate the overall single-cell transcriptome features of KD CAL and non-CAL patients, we collected the PBMCs from 3 CAL KD patients and 3 non-CAL KD patients. 3 CAL KD patients in this study were received IVIG treatment. The PBMCs of non-CAL KD patients were collected before and after IVIG treatment. Thus, there were three sample groups in our study: KD CAL patients (KD AT CAL), KD non-CAL patients before treatment (KD BT non-CAL), and KD non-CAL patients after treatment (KD AT non-CAL). After quality control and filter, the total number of detected cells was 77758, including 31530 cells from KD CAL patients, 26503 cells from KD AT non-CAL patients, and 19725 cells from KD BT non-CAL patients. To describe the scRNA-seq profiles in three PBMC groups, we first integrated nine samples together and clustered the cells across samples according to their expression features (Figure 1A and Supplemental Figure 1A). The detected cells could be classified into 19 major cell types including B cells, CD4T cells, CD8T cells, CD14 monocytes (CD14 mono), CD16 monocytes (CD16 mono), plasmacytoid dendritic cells (pDC), type 2 conventional dendritic cells (cDC2), erythrocytes and mixed cells (Eryth/mixed), gamma-delta cells and other T cells (gdT/other T), natural killer cells (NK), plasma blast cells (Plasmablast), platelets, hematopoietic stem and progenitor cells (HSPCs), mixed proliferating T cells (mixed proliferating T), mixed cells (mixed), and mixed T cells (mixed T). Both multimodal PBMC reference dataset and the canonical gene markers were used to validate major cell types (Supplemental Figure 1B and 1c).

**Figure 1.**
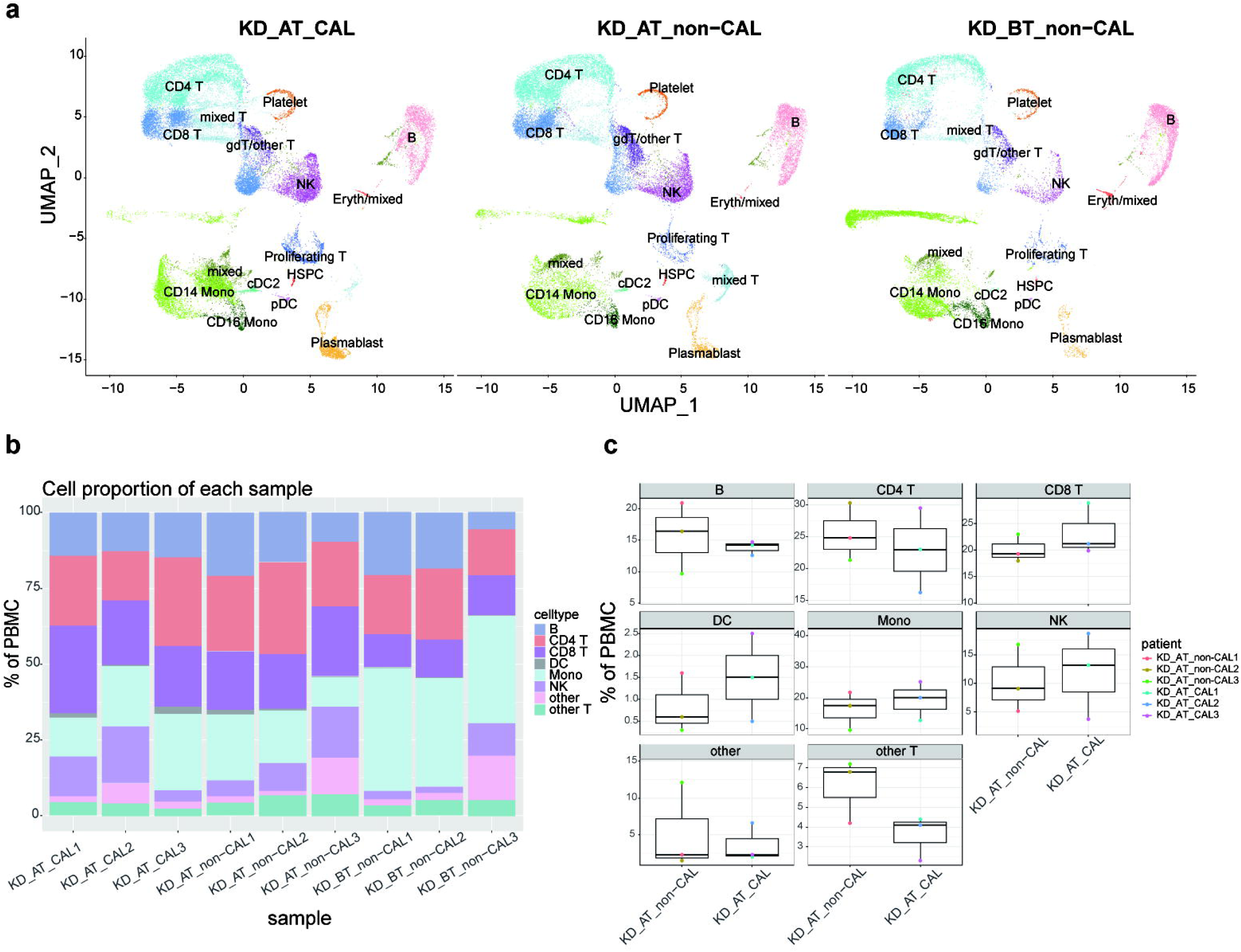
Single-cell profiling of PBMCs in KD CAL and non-CAL patients. **A.** The integration single-cell profiling analysis of nine samples from KD CAL patients after IVIG treatment (KD AT CAL), KD non-CAL patients after IVIG treatment (KD AT non-CAL), and KD non-CAL patients before IVIG treatment (KD BT non-CAL). **B.** The proportion of different cell types in each individual sample. **C.** Comparison of the proportion of major cell types between KD AT CAL and KD AT non-CAL patients.

### The proportions of different cell types between KD CAL patients and KD non-CAL patients after treatment

The percentages of major cell types for each patient are show in Figure 1b. Our scRNA-seq data demonstrated that KD patients after treatment had an increased percentage of plasma blast cells (p<0.05), T cells (CD4 T and CD8 T) and NK cells (p<0.05) and a decreased percentage of B cells and monocytes (p<0.05) compared with those before treatment, which were also well established by previous study(Supplemental Figure 2)^21^. We observed no such increased and decreased percentages of major cell types between KD AT CAL patients and KD AT non-CAL patients (Figure 1c). It proposes that IVIG treatment is at least effective for correcting the tilted major cell percentages in KD patients.

### Overall expression features of all single cells in KD CAL and non-CAL patients

We examined the expression features of all single cells in CAL KD and non-CAL KD patients. The differentially expressed genes (DEGs) were identified among two comparison groups: KD AT CAL patients vs. KD AT non-CAL patients, and KD AT CAL patients vs. KD BT non-CAL patients. The differentially expressed genes (DEGs) were identified in KD AT CAL patients using KD AT non-CAL and KD BT non-CAL patients as match groups (Figure 2a and 2b). When match group is KD AT non-CAL patients, there are 68 upregulated genes and 24 downregulated genes detected in KD AT CAL patients (p<0.05, Figure 2a). When match group is KD BT non-CAL patients, there are 168 upregulated genes and 159 downregulated genes detected in KD AT CAL patients (p<0.05, Figure 2b). Figure 2c shows that overlapped genes among four DEG groups. There are shared 11 downregulated genes in KD AT CAL patients compared to KD BT non-CAL and KD AT non-CAL patients. There are 63 shared upregulated genes in KD AT CAL patients compared to KD BT non-CAL and KD AT non-CAL patients. These DEGs actually reflect the expression difference between CAL and non-CAL patients. GO analysis shows that the shared upregulated genes in KD AT CAL patients are mainly involved in immunoglobulin and adaptive immunity while the shared downregulated genes in KD AT CAL patients are mainly involved in B cell activation and B cell receptor signaling pathway (Figure 2d and 2f). Thus, overall expression features of all single cells in KD AT CAL patients propose that they have a prolonged immune response and impaired B cell function compared to KD non-CAL patients.

**Figure 2.**
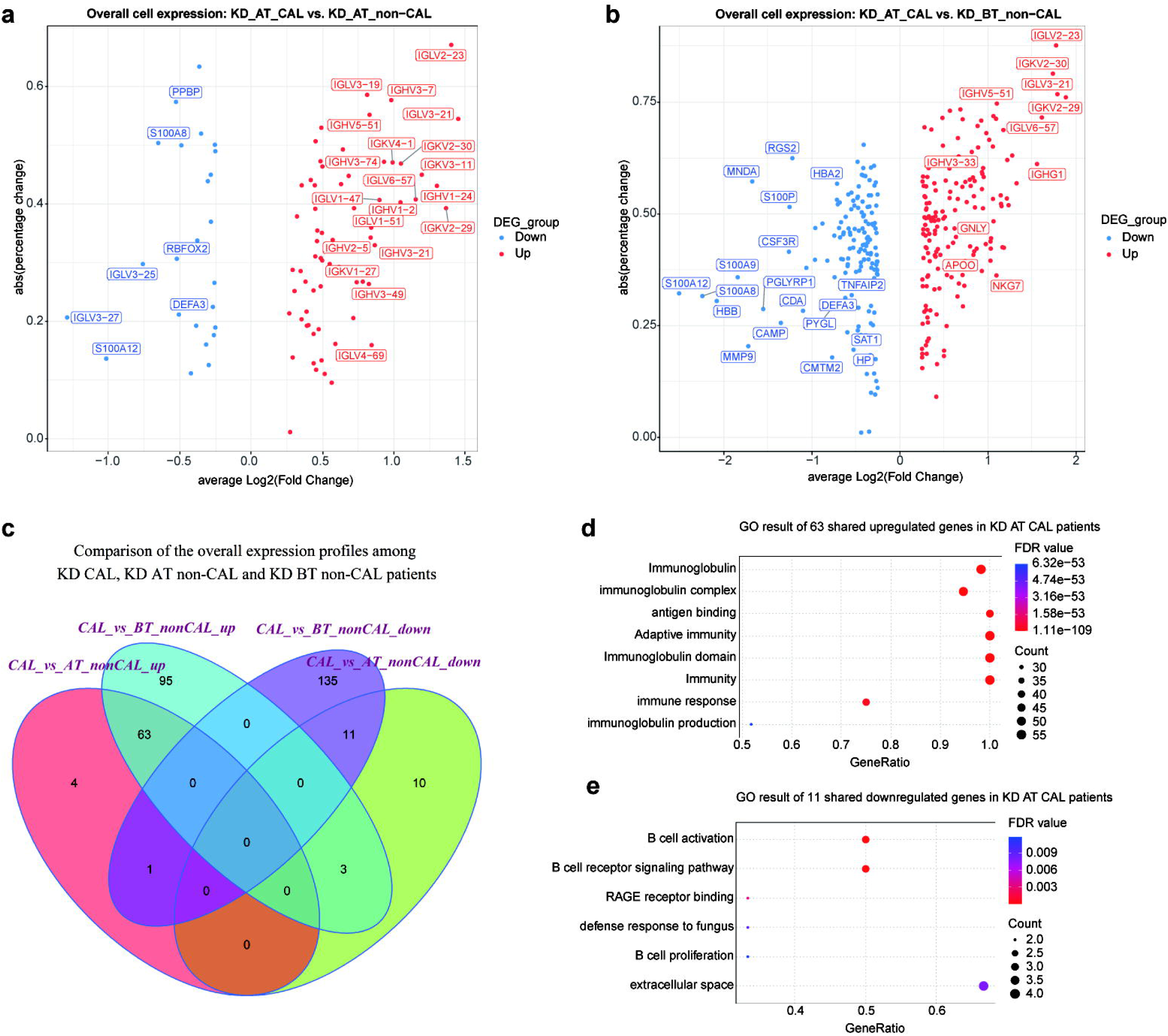
Expression analyses of all single-cells in KD CAL and non-CAL patients. A. Volcano plot of upregulated and downregulated genes in KD AT CAL patients compared to KD AT non-CAL patients (dots for only statistically significant genes and gene names for highest fold changes). B. Volcano plot of upregulated and downregulated genes in KD AT CAL patients compared to KD BT non-CAL patients (dots for only statistically significant genes and gene names for highest fold changes). C. Venn diagram of upregulated genes and downregulated genes for KD AT CAL patients in all four comparison groups. D. GO term enrichment analysis of 63 shared upregulated genes in KD AT CAL patients. E. GO term enrichment analysis of 11 shared downregulated genes in KD AT CAL patients.

### Pseudo-time analyses of PBMCs in KD CAL and non-CAL patients

Overall expression analysis shows that the gene related to B cell activation and B cell receptor signaling pathway are commonly repressed in KD AT CAL patients. It suggests that KD CAL patients may have malfunctional B cells even after IVIG treatment. The B cell development dysregulation has been studied in KD non-CAL patient^21^. Whether such phenomenon is present in KD AT CAL patients is worth investigation. We used pseudo-time analysis to reconstruct the cell developmental trajectory of PBMCs in KD AT CAL, KD AT non-CAL and KD BT non-CAL patients. The pseudo-time analyses show that the PBMCs in KD AT CAL patients also have three states which can be roughly classified as T/NK lineage (state 2), monocyte/B linage (state 1), and a shortened transitional state 3 (Figure 3a and 3b); the PBMCs in KD AT non-CAL patients have five states which can be roughly classified as B lineage (state 2), monocyte lineage (state 1), NK/platelet linage (state 4), T lineage (state 5), and a transitional state 3 (Figure 3c and 3d); the PBMCs in KD BT non-CAL patients have three states which can be roughly classified as monocyte/platelet lineage (state 1), T/B lineage (state 3), and monocyte lineage (state 2)(Figure 3e and 3f). These results demonstrates that KD AT non-CAL patients have a clearly differentiated cell developmental trajectory compared to KD BT non-CAL and KD AT CAL patients. For KD BT non-CAL patients, B cells and T cells failed to develop into two lineages. For KD AT CAL patients, B cells and monocytes failed to develop into two lineages. Thus, the poor differentiation problem for KD AT CAL patients is likely to be the developmental node between B cells and monocytes.

**Figure 3.**
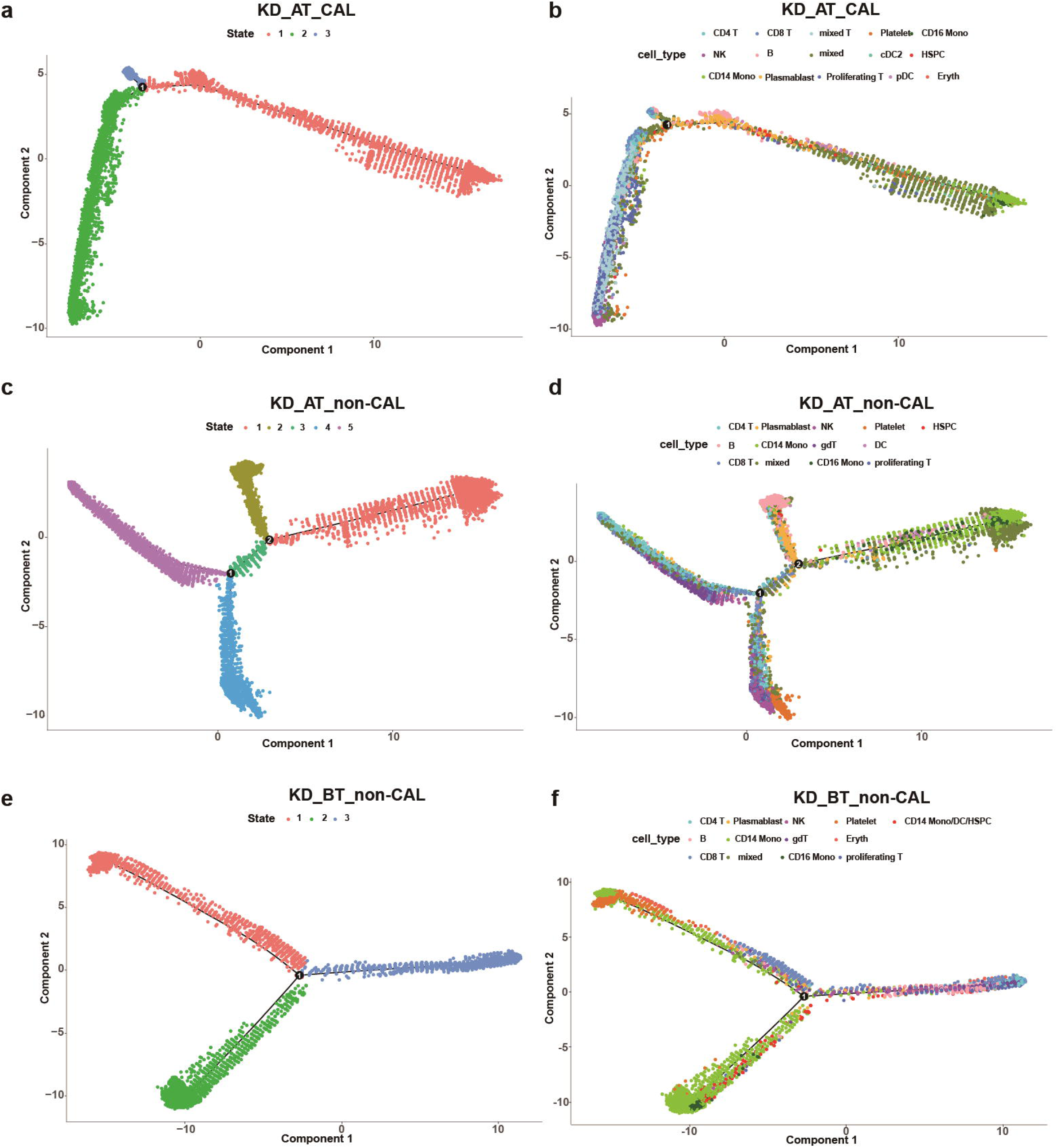
Pseudo-time analysis of cell developmental trajectory in KD CAL and non-CAL patients. A. The differentiation trajectory of all cells in KD AT CAL patients by states. B. The differentiation trajectory of all cells in KD AT CAL patients by cell types. C. The differentiation trajectory of all cells in KD AT non-CAL patients by states. D. The differentiation trajectory of all cells in KD AT non-CAL patients by cell types. E. The differentiation trajectory of all cells in KD BT non-CAL patients by states. F. The differentiation trajectory of all cells in KD BT non-CAL patients by cell types.

We further examined the expression dynamic of the genes related to cell cycle and B cell development in pseudo-time analyses. The expression dynamics of UBE2C, HSPD1, HSPE1, MT2A, MYC, and SPI1 were plotted along cell developmental trajectory based on pseudo-time analysis.UBE2C, HSPD1, HSPE1, MT2A, and MYC are the genes involved in cell cycle and SPI1 activates gene expression during B cell development^21^.We used HSPCs as the root for the cell developmental trajectory of each dataset, because they are the starting point of PBMCs. We found that there all six genes demonstrate a similar expression dynamic between KD AT CAL and KD AT non-CAL patients while they exhibit a different expression dynamic between KD BT non-CAL and KD AT non-CAL patients (Figure 4a, 4b and 4c). Both KD AT CAL and KD AT non-CAL patients received IVIG treatment which repressed the early expression of HSPD1, HSPE1, and MYC seen in KD BT non-CAL patients and restored the early expression of SPI1. The main expression dynamic differences between KD AT CAL and KD AT non-CAL patients are that SPI1 has a much higher expression slope in the early stage of cell development and MT2A has a much earlier expression peak in KD AT CAL patient s (Figure 4a and 4b).

**Figure 4.**
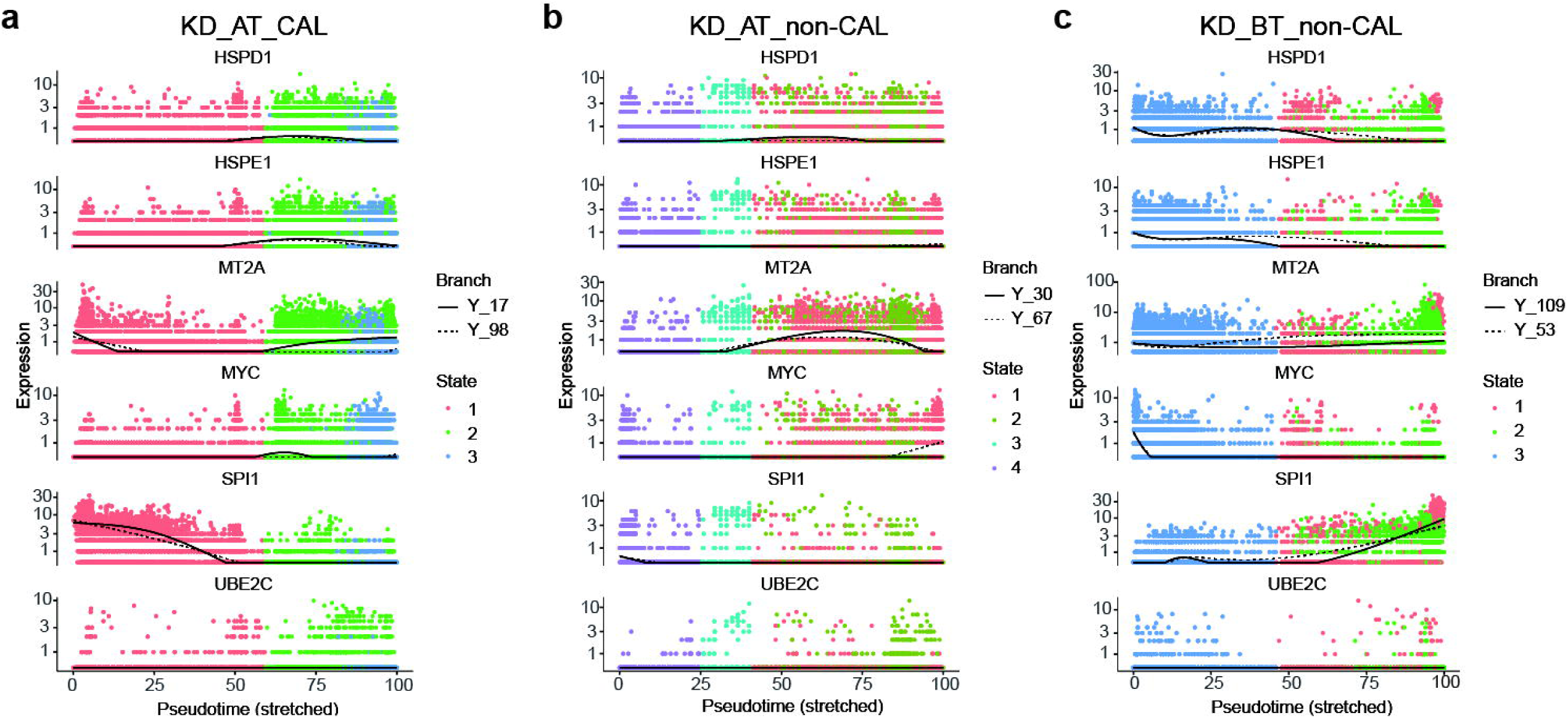
Pseudo-time analyses of expression dynamics of six B cell development and cell-cycle related genes in KD CAL and non-CAL patients. A. Expression dynamics of six B cell development and cell-cycle related genes in KD AT CAL patients. B. Expression dynamics of six B cell development and cell-cycle related genes in KD AT non-CAL patients. C. Expression dynamics of six B cell development and cell-cycle related genes in KD BT non-CAL patients.

### Cell communication patterns in KD CAL and non-CAL patients

We first calculated the total cell-to-cell interaction numbers and strength in in KD AT CAL, KD AT non-CAL and KD BT non-CAL patients. They provided an overview for cell communication setting for each sample group. Among three sample groups, the PBMCs of KD AT CAL patients have the intermediate number of interactions but the highest interaction strength; the PBMCs of KD AT non-CAL patients have the lowest number of interaction and the intermediate interaction strength while those of KD BT non-CAL patients have the highest number of interactions but the lowest interaction strength (Figure 5a). If using the signaling pattern in KD AT non-CAL patients as normal control, it can be seen that the intermediate number of interactions with the intermediate interaction strength is a recuperative pattern for KD patients. Thus, the problem with KD BT non-CAL patients is that they have a disordered signaling pattern with too many interactions of low strength, while the problem with KD AT CAL patients is that they must have some strong but bad signals that are related to CAL.

**Figure 5.**
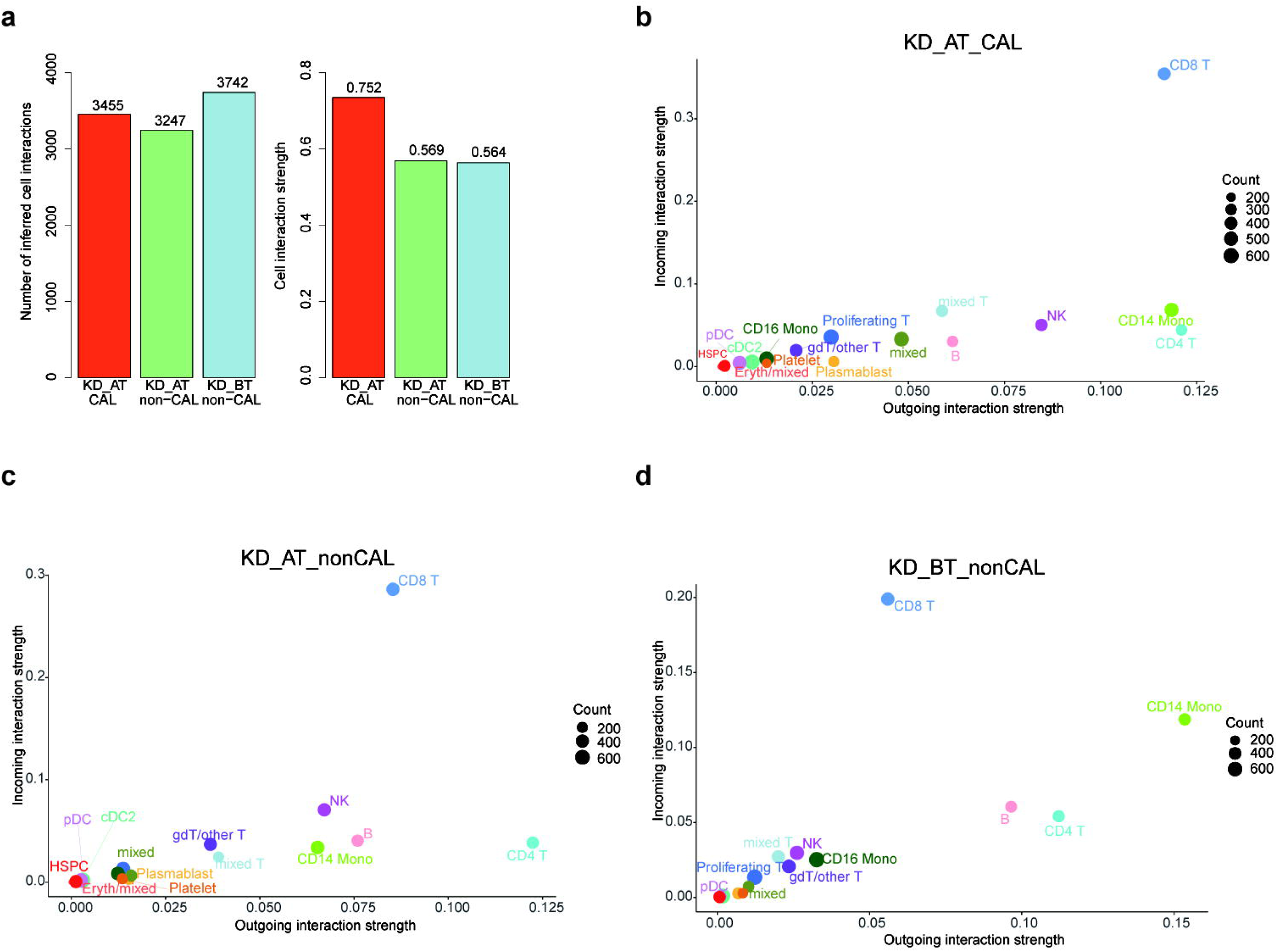
Cell-to-cell communication results in KD CAL and non-CAL patients. A. The number of cell-cell interactions and interaction strength in KD AT CAL, KD AT non-CAL, and KD BT non-CAL patients. B. The incoming and outgoing interaction strength of all cell types in KD AT CAL patients. C. The incoming and outgoing interaction strength of all cell types in KD AT non-CAL patients. D. The incoming and outgoing interaction strength of all cell types in KD BT non-CAL patients.

We further examined the signaling pattern among different types of cells in KD AT CAL, KD AT non-CAL and KD BT non-CAL patients. In KD AT CAL patients, cDC2 cells have the highest number of outgoing signals and CD16 monocytes have the highest number of incoming signals (SupplementalFigure3a); CD4 T and CD8 T cells have the highest outgoing and incoming signal strength as in KD AT non-CAL patients (Figure 5b). In KD AT non-CAL patients, cDC2 cells have the highest numbers of incoming and outgoing signals among all cell types (Supplemental Figure3b); CD4 T and CD8 T cells have the highest outgoing and incoming signal strength among all cell types, respectively (Figure 5c). In KD BT non-CAL patients, CD16 monocytes have the highest number of outgoing signals and cDC2 cells have the highest number of incoming signals (SupplementalFigure3c); CD8 T cells have the highest incoming signal strength and CD14 monocytes have the highest outgoing signal strength (Figure 5d). In both KD AT CAL and KD BT non-CAL patients, CD14 monocytes have the most significant change of outgoing signal strength compared to those in KD AT non-CAL patients. Because KD AT non-CAL patients have the highest interaction strength among three sample groups, it is inferred that the signal molecules in their CD14 monocytes might contribute to CAL development.

### Bad and good signaling molecules from CD 14 monocytes

In order to study the significant signaling difference in KD AT CAL patients, we compared the signaling changes of their monocytes to those of KD AT non-CAL and KD BT non-CAL patients (SupplementalFigure4a, 4b, 4c and 4d). Four signaling molecules from monocytes have distinctively contrasting expression patterns between KD CAL and non-CAL patients. The expression of APP, CCL, and MCH-IIsignal molecules is significantly increased in the CD14 monocytes of KD CAL patients while the expression of RESISTIN is significantly increased in those KD non-CAL patients (Figure 6a, 6b, 6c and 6d). These results propose that APP, CCL, MCH-I and MCH-II are bad indicators for KD patients for their relation with CAL while RESISTIN is a good one for KD patient. We further investigated upregulated and downregulated signaling ligand-receptor pairs in KD AT CAL patients (Supplemental Figure 5a, 5b, 5c and 5d). Compared to KD AT non-CAL and KD BT non-CAL patients, the HLA ligands from CD 14 monocytes to CD4 and CD8B receptors on T cells are upregulated in KD AT CAL patients while the RETN ligand from CD 14 monocytes to CAP1 receptors on all cell types is downregulated in KD AT CAL patients.

**Figure 6.**
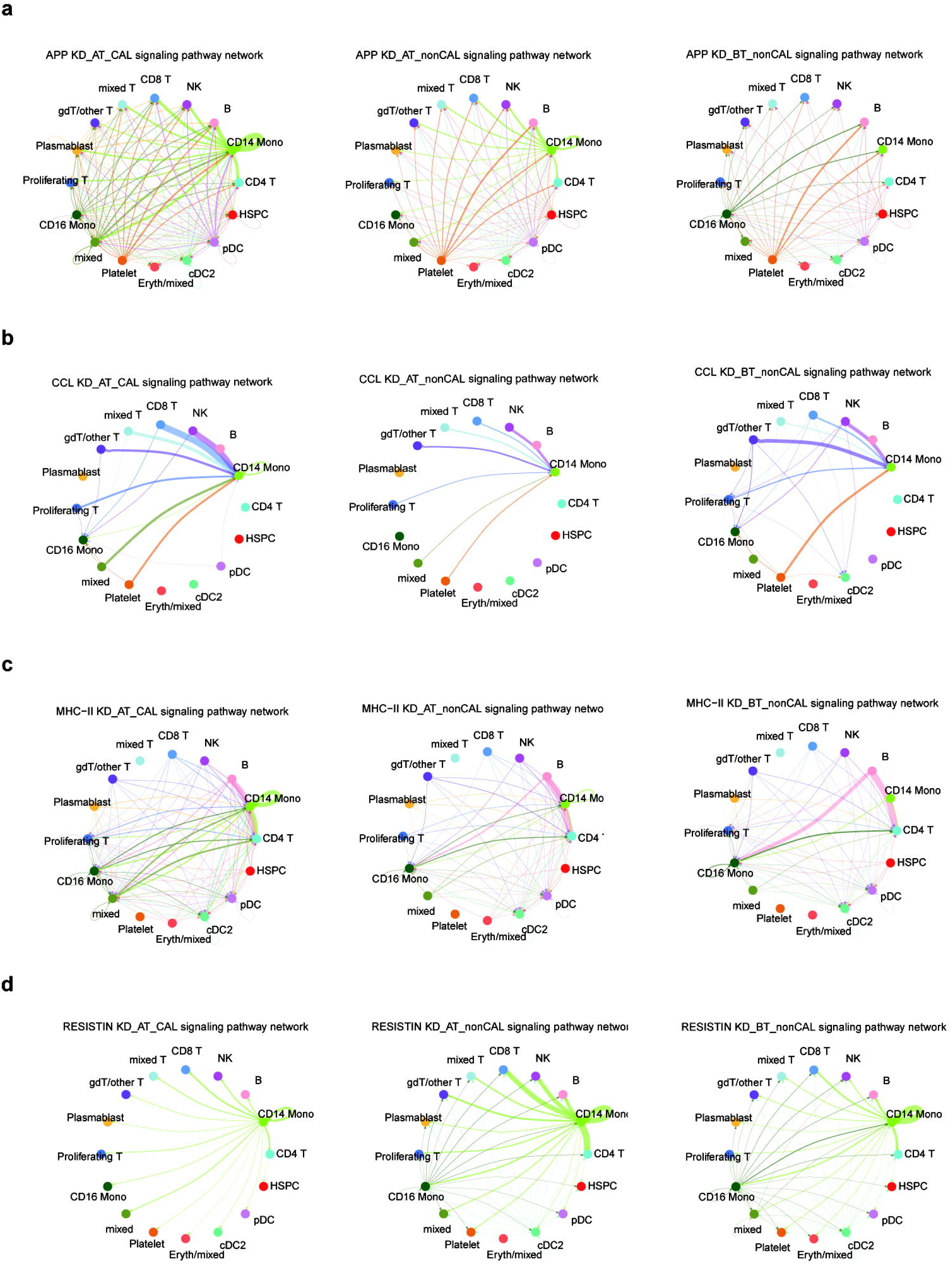
The signaling pathway network of four molecules in KD CAL and non-CAL patients. A. The APP signaling pathway network of all cell types in KD AT CAL, KD AT non-CAL, and KD BT non-CAL patients. B. The CCL signaling pathway network of all cell types in KD AT CAL, KD AT non-CAL, and KD BT non-CAL patients. C. The MCH II signaling pathway network of all cell types in KD AT CAL, KD AT non-CAL, and KD BT non-CAL patients. D. The RETN signaling pathway network of all cell types in KD AT CAL, KD AT non-CAL, and KD BT non-CAL patients. Note: the width of strip indicates the strength of signaling pathway.

The expression profile of CD 14 monocytes in KD AT CAL patients were also examined by comparing to that of KD AT non-CAL or KD BT non-CAL patients. Using KD AT non-CAL patients as control, there are 33 upregulated genes and 73 downregulated genes detected in KD AT CAL patients. Using KD BT non-CAL patients as control, there are 173 upregulated genes and 112 downregulated genes detected in KD AT CAL patients. There are 33 shared upregulated genes and 50 shared downregulated genes in both comparison groups (Figure 7a and 7b). GO analyses of these shared DEGs in KD AT CAL patients show that the shared upregulated genes are mainly involved in antigen processing and presentation and the shared downregulated genes are mainly involved in antimicrobial function and innate immunity ((Figure 7c and 7d). Antigen processing and presentation by MHC II signaling pathway is an important adaptive immune response. It explains why the overall expression features of KD AT CAL patients’ PBMCs exhibit an elevated level of adaptive immunity whose source is CD 14 monocytes.

**Figure 7.**
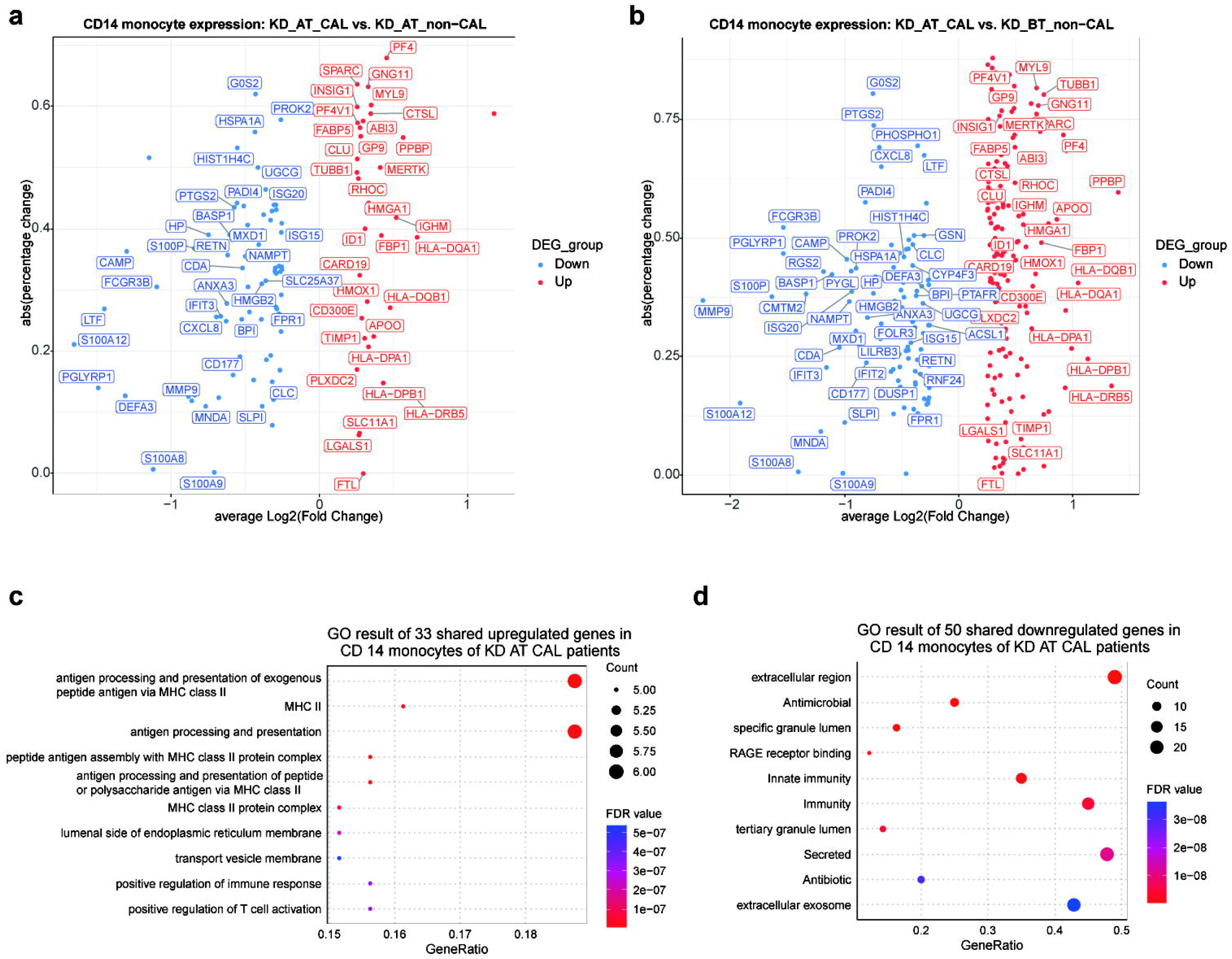
Expression analyses of CD14 monocytes in KD CAL and non-CAL patients. A. Volcano plot of upregulated and downregulated genes in the CD14 monocytes of KD AT CAL patients compared to those of KD AT non-CAL patients (dots for only statistically significant genes and gene names for the shared genes with KD BT non-CAL patients). B. Volcano plot of upregulated and downregulated genes in in the CD14 monocytes of KD AT CAL patients compared to those of KD BT non-CAL patients (dots for only statistically significant genes and gene names for the shared genes with KD AT non-CAL patients). C. GO term enrichment analysis of 33 shared upregulated genes in the CD14 monocytes of KD AT CAL patients. D. GO term enrichment analysis of 50 shared downregulated genes in the CD14 monocytes of KD AT CAL patients.

## Discussion

KD was first reported in 1967 and its etiology remains unclear. Our previous research shows that B cell developmental dysfunction is one of cellular contributions for KD development^21^. The over expression of HSPD1 and HSPE1 genes and the under expression of MYC, SPI1, MT2A and UBE2C genes are observed in the early cell development stage of KD patients before treatment. In this study, we mainly focused on the cellular features of KD AT CAL patients. After IVIG treatment, the expression of HSPD1 and HSPE1 genes was repressed while the expression of SPI1 and MT2A genes was rescued in KD AT CAL patients. The similar outcome could be seen in KD AT non-CAL patients as well. However, the expression pattern of SPI1 and MT2A in KD AT CAL patients are still different from that of KD AT non-CAL patients. Their early high expression explains the poor differentiation between B cell and monocytes in KD AT CAL patients. It further leads to the abnormal immune responses of CD 14 monocytes in CAL patients.

Further analysis of cell-to-cell communication patterns show that the expression level of C-C motif chemokine ligand (CCL) is significantly higher in the CD14 monocytes of KD AT CAL patients than in those who had not developed CALs. The actual upregulated signaling is CCL5-CCR1 ligand-receptor pair in KD AT CAL patients (Supplemental Figure 5a and 5c). CCL5 is a member of CC chemokine ligand family (CCL). The human CCL5 gene is located on chromosomes 17q11.2-q12 and consists of two introns and three exons. CCL5 composes of 68 amino acids and is expressed on various cell types, including T lymphocytes, macrophages, synovial fibroblasts, tubular epithelial cells and platelets. This chemokine, in combination with its receptor, CCR5, plays a variety of biological functions^22,23^ It has chemotactic or stimulating effect on white blood cells such as T cells, NK cells and monocytes. It causes the migration of lymphocytes and mediates the occurrence and reaction process of inflammation. It is also involved in the pathogenesis and development of many diseases such as HIV infection, cancer, and autoimmune disease. There is evidence of endothelial cell activation through CCL5^24,25^. A previous study based on 173 KD samples and 101 normal samples using differential analysis, weighted gene co-expression network analysis and co-expression network construction found that the CCL5 and four other genes as the key genes mediating the development of KD^26^. Furthermore, it has been reported that CCL5-403A variant may be associated with the coronary involvement in North Indian children with KD^27^. Like in the mentioned studies, our results suggest that CCL5 may be a potential biomarker of CALs in KD.

Cell-to-cell communication analyses also indicate that the gene amyloid beta precursor protein (APP) was upregulated in the CD14 monocytes of KD AT CAL patients compared to KD AT non-CAL and KD BT non-CAL patients. APP is deemed to be ubiquitously expressed, with high levels in the brain and low levels in endocrine tissue, bone marrow, liver, gall bladder, and in male and female reproductive organs, whose abnormal processing and/or overexpression is the basis of the amyloid β peptide (Aβ) found in the brains of patients with Alzheimer disease^28^. There is evidence that Aβ is involved in vascular degeneration and dysfunction^29^. Aβ can interact with fibrinogen to form plasmin-resistant abnormal blood clots^30^. Another study reported the direct evidence that endothelial cells are specifically susceptible to intracellular overexpression of APP, and that free radical generation is the likely mechanism of cell damage due to APP overexpression^31^. Although no previous studies have reported the involvement of APP in KD or CALs, considering its damage to endothelial cells, we suggest that it might be a key molecule in the development of CALs in KD patients.

The over expression of MHC II genes is expected in KD AT CAL patients, since the overall cell expression features show that they have an elevated level of adaptive immunity. Antigen processing and presentation which is mainly administrated by MHC II genesis an important component of adaptive immunity^32^. Several MHC-II molecules including HLA-DRB5, HLA-DRB1, and HLA-DRA are upregulated in the CD14 monocytes of KD AT CAL patients and they all bind to CD4 receptor (Supplemental Figure 5a and 5c). It proposes that the CD14 monocyte of KD AT CAL patients interact with their CD4 T cells and further magnify adaptive immune cascade by manipulating T cell receptor signaling pathway. This viewpoint is supported by the upregulation of MHC I genes in the CD14 monocyte of KD AT CAL patients. Several MHC-I molecules including HLA-A, HLA-B, and HLA-C are upregulated in the CD14 monocytes of KD AT CAL patients as well. They all bind to CD8A receptor, which indicates that CD8 T cells would respond to the MHC-I signals from CD14 monocyte. Thus, both CD4 and CD8 T cells are likely to take part in the prolonged adaptive immune response by the summons of the CD14 monocytes in KD AT CAL patients and play their part in the development of CALs.

RETN expression is a reverse indicator of CALs for KD patients revealed by cell-to-cell communication analyses. The outgoing signal strength of RETN from the CD14 monocytes of KD AT CAL patients is much weaker than that of KD AT non-CAL and KD BT non-CAL patients and the feedback signals of RETN in the other cell types almost disappear in KD AT CAL patients (Figure 6d). Using KD AT non-CAL or KD BT non-CAL patients as background, the downregulation of RETN-CAP1 signaling is present in all CD14 monocyte’s interactions to the other major cell types, especially to CD4 T, B, NK, and CD8 T cells (Supplemental Figure 5b and 5d). There is evidence that RETN has antibacterial activity which is a function of innate immunity^33^. The repression of RETN-CAP1 signaling is thus a sign of the downregulation of innate immunity, which is consistent with the overall cell expression features of KD AT CAL patients that B cell activation and B cell related biological processes are downregulated in them, because B cell plays an essential role in innate immunity.

This study has a limited number of CAL samples which might not provide a comprehensive explanation for the development of CALs in KD patients. We could not collect the sample of KD CAL before IVIG treatment. First, there is only a small proportion of KD patients who would develop CALs. We don’t know which patient would develop CALs before IVIG treatment. CAL development in KD patients usually occur after IVIG treatment, around 5-7 days of the disease course. However, in clinical practice, the diagnosis of KD is often confirmed between days 4-5 of the illness. Therefore, obtaining samples of KD combined with CALs before IVIG treatment is difficult. Here we propose to establish a CAL cohort from multiple centers in future research in order to collect KD CAL patients before and after IVIG treatment, which would provide a better understanding for the pathogenesis of CALs in KD patients.

In conclusion, we delineated the single-cell atlas of PBMCs for KD AT CAL, KD AT non-CAL and KD BT non-CAL patients in this study. Our results demonstrate that KD CAL patients have a prolonged adaptive immune response and repression of B cell function related genes’ expression after IVIG treatment. It further leads to the poor cell differentiation between B cell lineage and monocyte lineage. Cell-to-cell communication analyses further show that CD14 monocytes have the most significant change of outgoing signal strength in all major cell types of KD AT CAL patients compared to those of KD AT non-CAL patients. The further analyses discover four differentially expressed prominent signaling molecules in the CD14 monocytes of KD AT CAL patients which are APP, CCL, MCH-II, and RETN. APP, CCL, and MCH-II are upregulated signaling molecules in the CD14 monocytes of KD AT CAL patients while RTEN is downregulated one. The expression profile of the CD14 monocytes of KD AT CAL patients exhibits a pattern of high adaptive immune response and low innate immune response. The results above suggest that CD14 monocytes might be one of major cellular contributors for CAL development in KD patients. They first express a hyper adaptive immune tendency and then manipulate CD4 and CD8 T cells to join the same process. Along the process, the normal function of B cells is suppressed and several key signaling molecules are either upregulated or downregulated to further trigger CAL onset in KD CAL patients who have a persistent immune response compared to non-CAL patients. Among key signaling molecules, APP, CCL, and MCH-II could serve as the indicator for CAL development and RETN could serve as the reverse indicator CAL development. Especially, APP and CCL are two genes associated with coronary involvement and damage to endothelial cells.

## Materials and Methods

### Participants and clinical characteristics

All participants were recruited from Shanghai Children’s Hospital. All manipulations were approved by the Ethics Committee of Shanghai Children’s Hospital (IRB number: 2022R121). Guardians of the participants had provided their informed consent in the study. KD was diagnosed according to the diagnosis criteria established by the American Heart Association^1^. KD patients were collected blood samples 24 hours after IVIG therapy and subsidence of fever.

#### 1. Single-cell RNA sequencing and data analyses

##### 2.1 scRNA-seq library construction

Peripheral blood samples (2 mL each sample) were collected from the participants. PBMCs of participants were isolated according to standard density gradient centrifugation methods by using the Ficoll-Paque medium. The cell viability should exceed 90%. The single-cell library was constructed using the 5’ Library Kits. The cell suspension was loaded onto a chromium single-cell controller (10X Genomics) to generate single-cell gel beads in the emulsion (GEMs) according to the manufacturer’s protocol. Lysis and barcoded reverse transcription of polyadenylated mRNA from single cells was performed inside each GEM. Post-RT-GEMs were cleaned up, and cDNA was amplified. The barcoded sequencing libraries were generated using the Chromium Next GEM Single Cell V(D)J Reagent Kits v1.1 (10x Genomics) and were sequenced as 2×150-bp paired-end reads on an Illumina NovaSeq platform.

##### 2.2 scRNA-seq data processing

Cell Ranger (Version6.0.0) software was used to process the raw FASTQ files, align the sequencing reads to the GRCh38 reference transcriptome and generate a filtered UMI expression profile for each cell. Raw gene expression matrices were read into R (Version 4.2.1) and converted to Seurat objects. The number of genes, UMI counts and percentage of mitochondrial genes were examined to identify outliers. The following criteria was applied for quality control: total UMI count between 2,000 and 60,000, and mitochondrial gene percentage <5%. After removal of low-quality cells, the count matrix was normalized by SCTransform method, which is based on a negative binomial regression model with regularized parameters. Then all datasets from the individual samples were integrated using the “Find Integration Anchors” and “Integrate Data” functions in Seurat (Version 4.1.1)^34^. We identified “anchors” among all individual datasets with multiple canonical correlation analysis (CCA) and used these “anchors” to create a batch-corrected expression matrix of all cells, which allowed the cells from different datasets to be integrated and analyzed. The supervised principal component analysis (SPCA) was performed to reduction and the weighted nearest neighbor(wnn) graph-based clustering was used to identify cell clusters. The cell identities were determined with multimodal reference mapping in Seurat (Version 4.1.1)^34^.

##### 2.3 Differential expression and functional enrichment analysis

DEG analysis for each cell type on the sample level following the recommendation of Bioconductor^35^. The differential expression analysis was conducted between conditions by using Libra (Version 1.0.0)^36^, which implements a total of 22 unique differential expression methods that can all be accessed from one function. We used “run_de” functions with pseudo-bulk approach, implementing the DESeq2 (Version 1.36.0)^37^ with a likelihood ratio test. GSEA pathway analysis was performed using cluster Profiler (Version 4.7.1)^38^. Hallmark gene sets in the Molecular Signatures Database (MSigDB Version7.1)^39^ served as the gene function database. P-values were adjusted to FDRs. FDRs < 0.05 was chosen as the cut-off criterion indicating a statistically significant difference.

##### 2.4 Pseudo-time analysis of cell differentiation trajectories

Pseudo-time analysis of cell differentiation trajectories for each sample dataset was performed with R package Monocle 2^40^. The expression feature and inferred cell type for each sample dataset from Seurat result was used to construct the cell dataset for Monocle analysis pipeline. We used the Monocle built-in approach named “dpFeature” to detect the variable genes that define cell’s differentiation. Its advantages are needing no prior biological knowledge and discovering important ordering genes from data itself. Dimension reduction was performed with 2 max components and “DDR Tree” method. HSPCs and B cells in each sample dataset for pseudo-time analysis were extracted according to the cell identities from Seurat result.

##### 2.5 Cell-cell communication analysis

Cell-cell communication analysis was performed with R package CellChat 1.6.1^41^. Human database in CellChat was set to be the default ligand-receptor pair for cell-to-cell communication analysis.

## Acknowledgements

We would like to thank all the donors who contributed samples.

## Author contributions

L.X., L.S. and T.X. conceived and designed the study. X.L., Q.Q, S.S., L.C., X.J., W.L. and M.H. contributed to collected samples and clinical information. Q.L., X.L., Q.Q. and L.S. performed bioinformatic analyses and discussed and interpreted the data. L.X., L.S, Q.L. and L.M. wrote the manuscript. L.X. and T.X. revised it.

## Informed consent

The study protocol was approved by the Ethics Committee of Shanghai Children’s Hospital (IRB number: 2022R121). Guardians of the participants had provided their informed consent in the study.

## Data availability statement

The raw sequence data are available under restricted access because of data privacy laws, and access can be obtained by reasonable request to the corresponding authors.

