## Supplementary figures and images for "Delineation of Single-cell Altas Provides New Insights for Development of Coronary Artery Lesions in Kawasaki Disease: Bad and Good Signaling Molecules"

### Supplemental Figure1

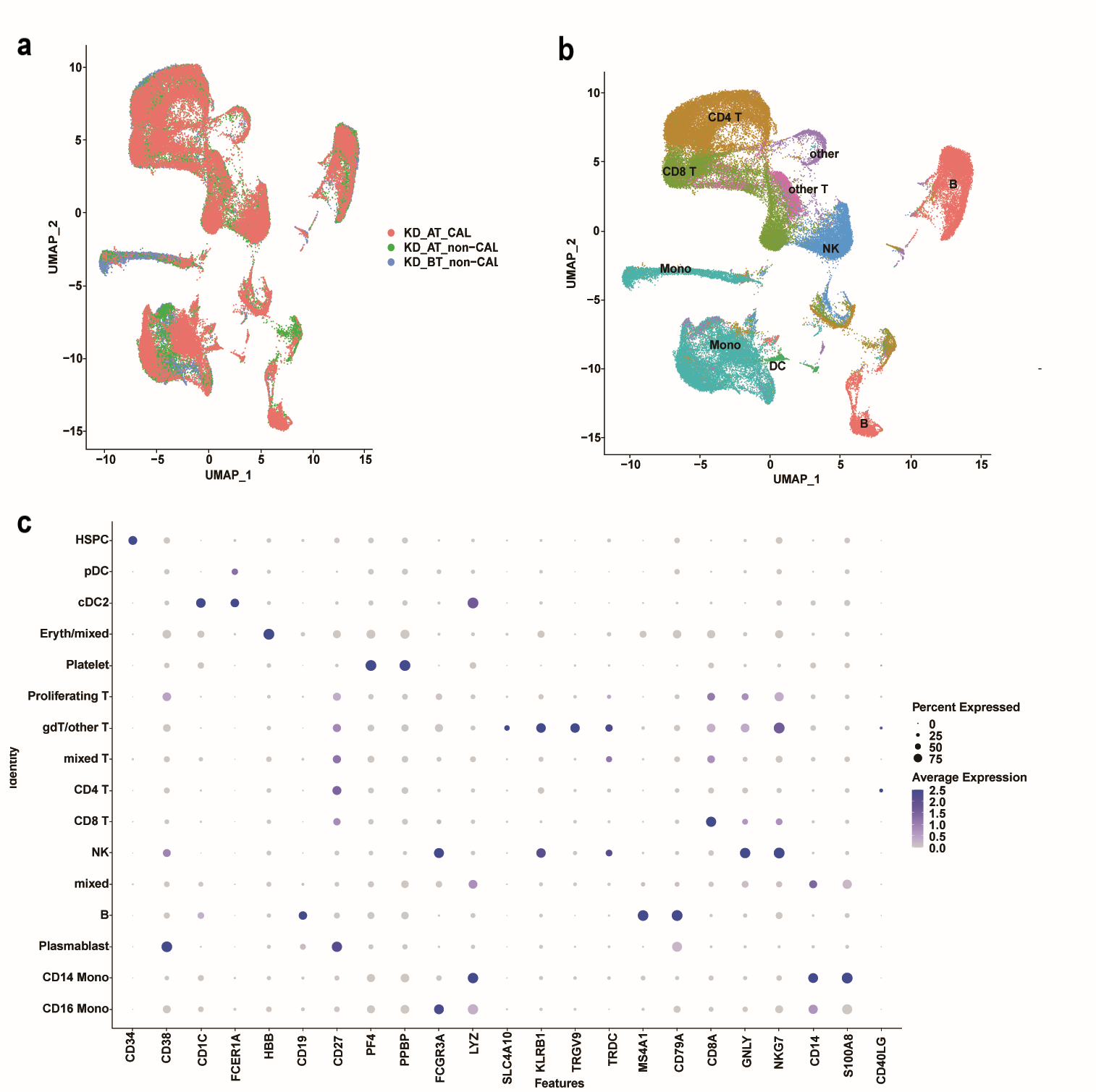

### Supplemental Figure2

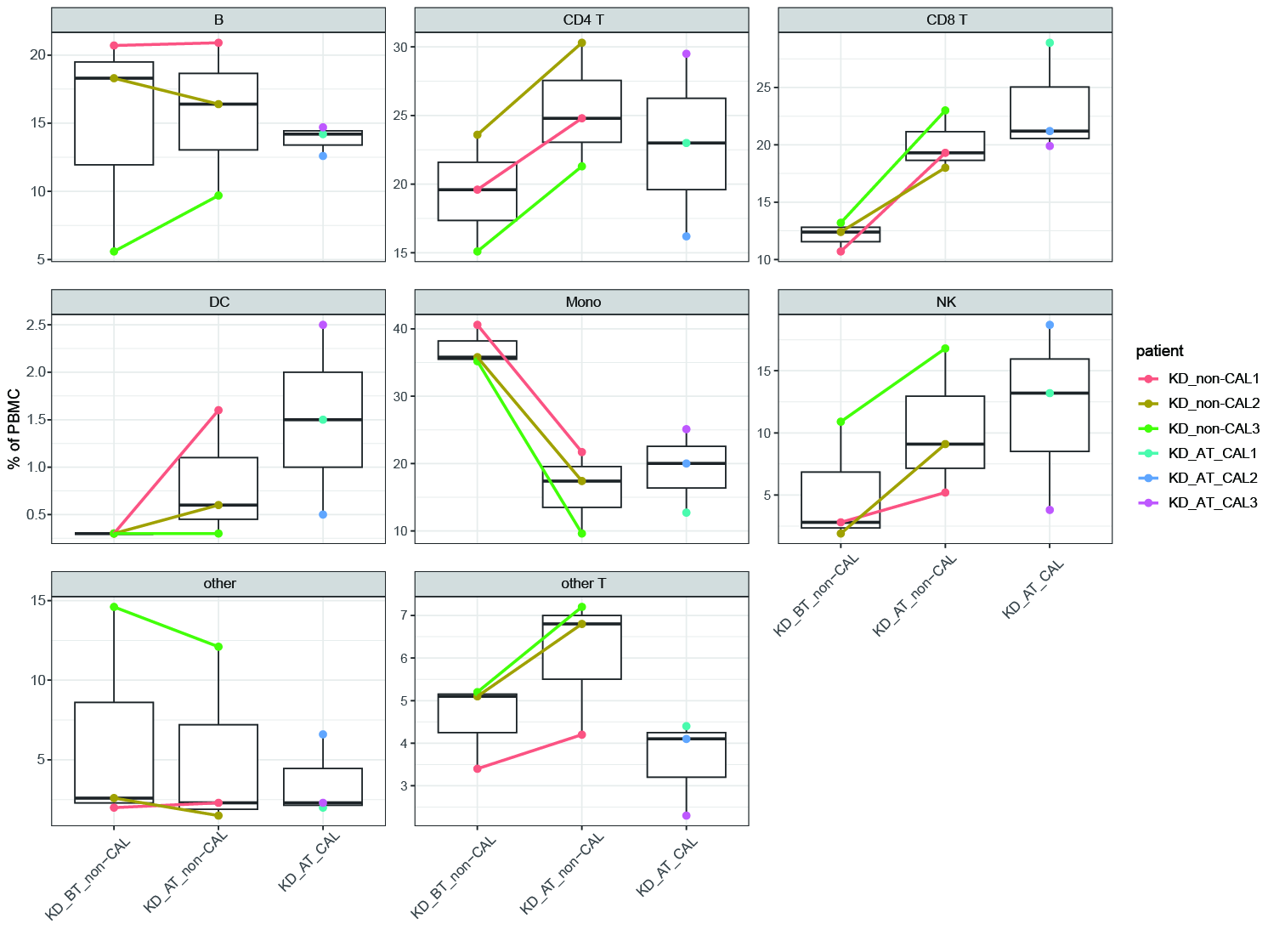

### Supplemental Figure3

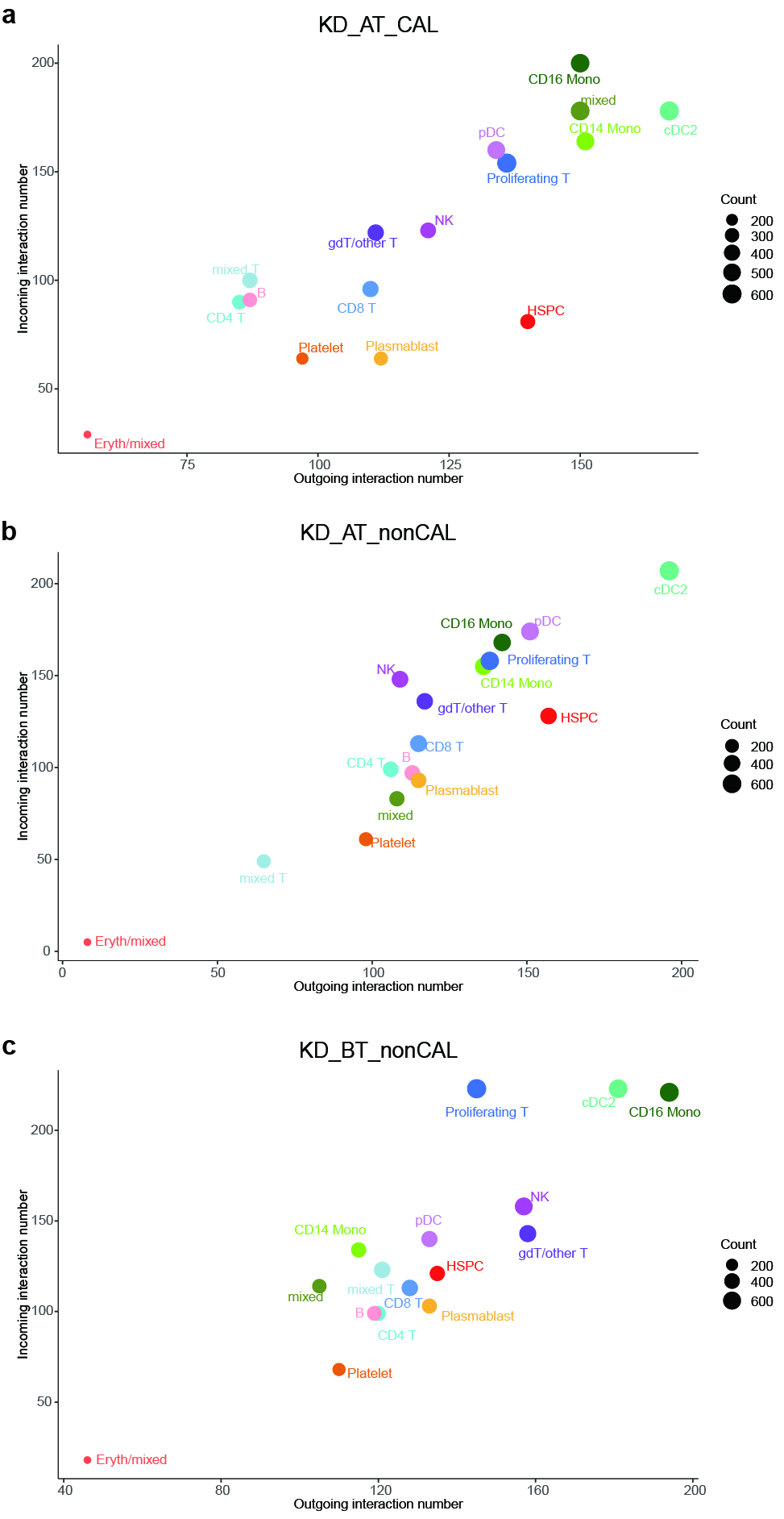

### Supplemental Figure4

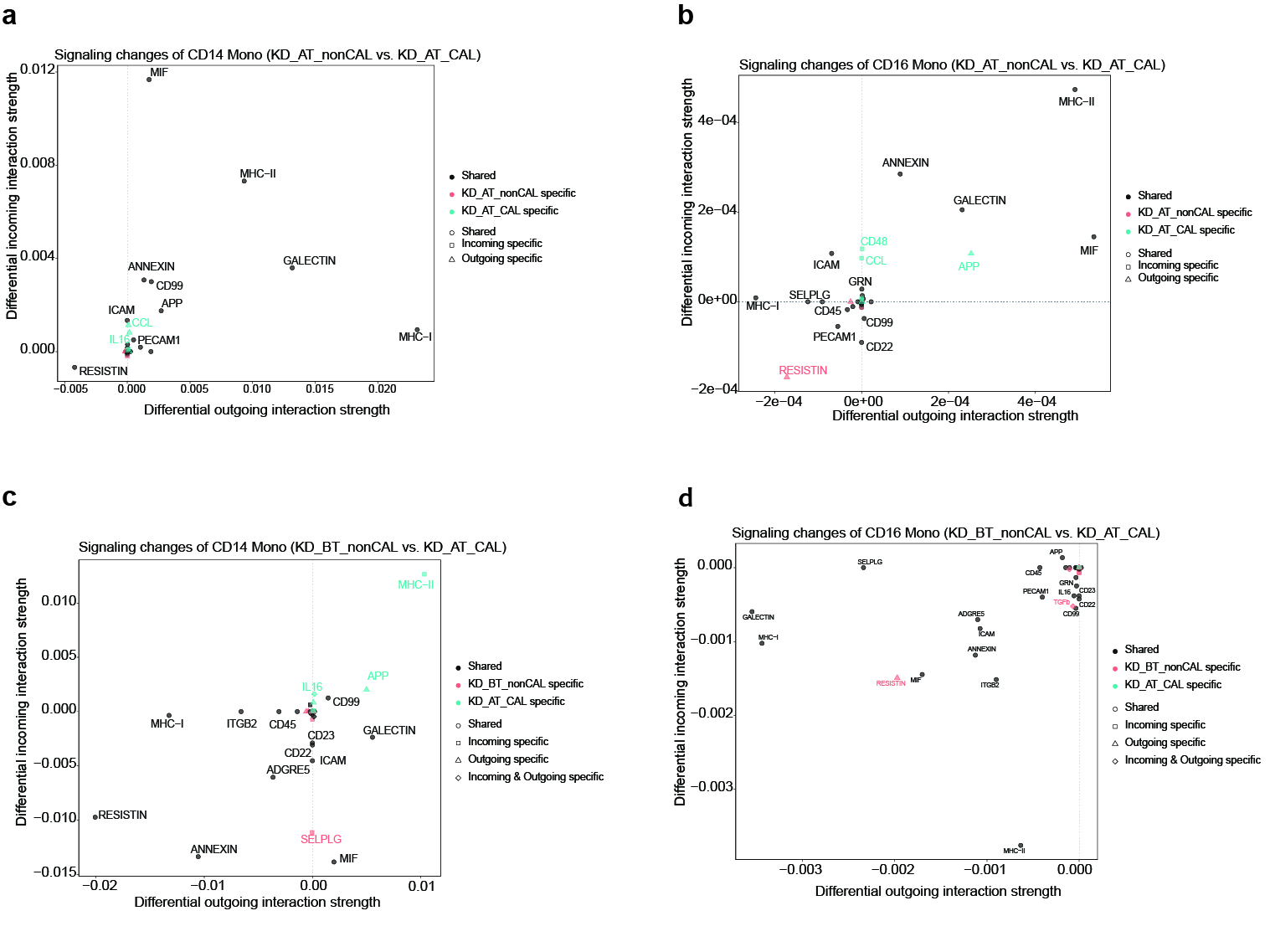

### Supplemental Figure5

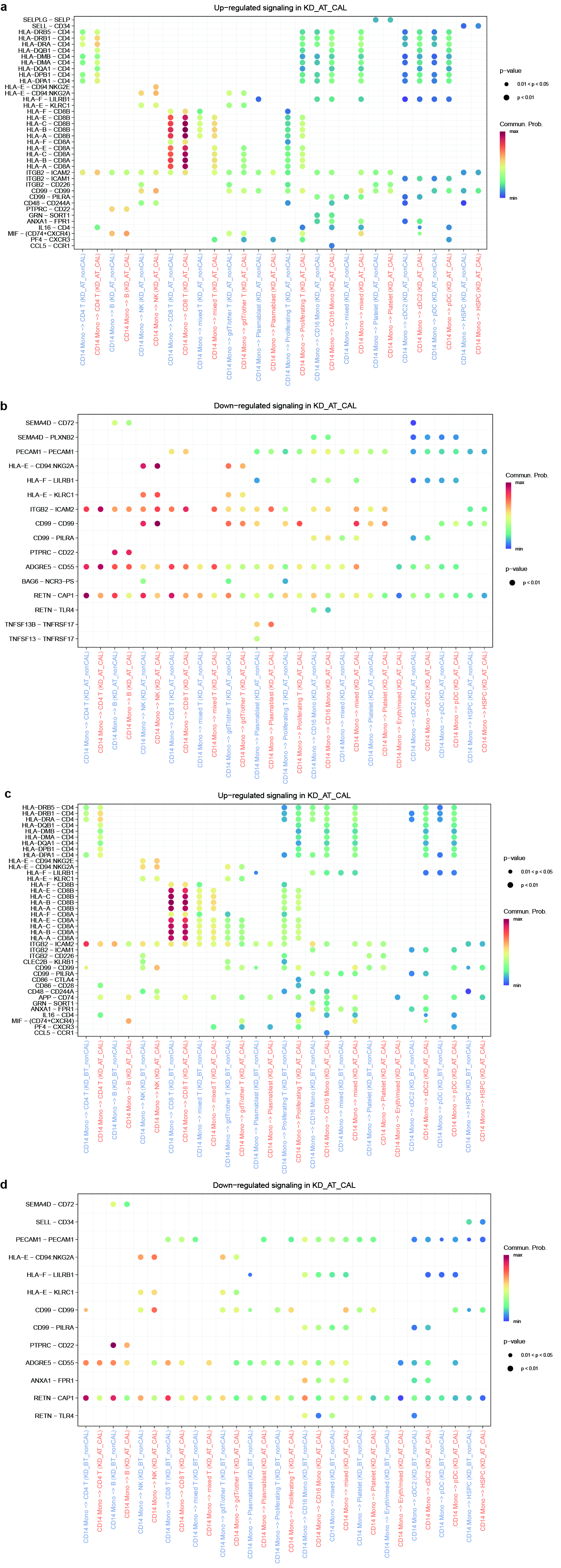
