## Supplemental Figure legends for "Delineation of Single-cell Altas Provides New Insights for Development of Coronary Artery Lesions in Kawasaki Disease: Bad and Good Signaling Molecules"

**Supplemental figure 1.** Integrated single-cell profiling of PBMCs in KD CAL and non-CAL patients. A. Integrated single-cell profiling of PBMCs in KD CAL and non-CAL patients. B. The inferred cell types marked with different colors based on multimodal reference mapping. C. Expression of canonical gene markers for each cell type in KD CAL and non-CAL patients based on integration analysis.

**Supplemental figure 2.** Comparison of the proportion of major cell types among KD AT CAL, KD AT non-CAL patients and KD BT non-CAL patients.

**Supplemental figure 3.** Cell-to-cell communication results in KD CAL and non-CAL patients. A. The incoming and outgoing interaction numbers of all cell types in KD AT CAL patients. B. The incoming and outgoing interaction numbers of all cell types in KD AT non-CAL patients. C. The incoming and outgoing interaction numbers of all cell types in KD BT non-CAL patients.

**Supplemental figure 4.** Signaling changes of CD14 and CD16 monocytes in KD AT CAL patients. A. Signaling changes of CD14 monocytes in KD AT CAL patients compared to those in KD AT non-CAL patients. B. Signaling changes of CD16 monocytes in KD AT CAL patients compared to those in KD AT non-CAL patients. C. Signaling changes of CD14 monocytes in KD AT CAL patients compared to those in KD BT non-CAL patients. D. Signaling changes of CD16 monocytes in KD AT CAL patients compared to those in KD BT non-CAL patients.

**Supplemental figure 5.** Upregulated and downregulated signaling of CD14 monocytes in KD AT CAL patients. A. Upregulated signaling of CD14 monocytes in KD AT CAL patients to the other cell types in both KD AT CAL patients and KD AT non-CAL patients. B. Downregulated signaling of CD14 monocytes in KD AT CAL patients to the other cell types in both KD AT CAL patients and KD AT non-CAL patients. C. Upregulated signaling of CD14 monocytes in KD AT CAL patients to the other cell types in both KD AT CAL patients and KD BT non-CAL patients. D. Downregulated signaling of CD14 monocytes in KD AT CAL patients to the other cell types in both KD AT CAL patients and KD BT non-CAL patients.
